# Computational modeling of macrophage iron sequestration during host defense against *Aspergillus*

**DOI:** 10.1101/2022.01.24.477648

**Authors:** Bandita Adhikari, Yogesh Scindia, Luis Sordo Vieira, Henrique de Assis Lopes Ribeiro, Joseph Masison, Ning Yang, Luis L. Fonseca, Matthew Wheeler, Adam C. Knapp, Yu Mei, Brian Helba, Carl Atkinson, Will Schroeder, Borna Mehrad, Reinhard Laubenbacher

## Abstract

Iron is essential to the virulence of *Aspergillus* species, and restricting iron availability is a critical mechanism of antimicrobial host defense. Macrophages recruited to the site of infection are at the crux of this process, employing multiple intersecting mechanisms to orchestrate iron sequestration from pathogens. To gain an integrated understanding of how this is achieved in invasive aspergillosis, we generated a transcriptomic time-series of the response of human monocyte-derived macrophages to *Aspergillus* and used this and the available literature to construct a mechanistic computational model of iron handling of macrophages during this infection. We found an overwhelming macrophage response beginning 2-4 hours after exposure to the fungus, which included upregulated transcription of iron import proteins transferrin receptor-1, divalent metal transporter-1, and ZIP family transporters, and downregulated transcription of the iron exporter ferroportin. The computational model, based on a discrete dynamical systems framework, consisted of 21 3-state nodes, and was validated with additional experimental data that were not used in model generation. The model accurately captures the steady state and the trajectories of most of the quantitatively measured nodes. In the experimental data, we surprisingly found that transferrin receptor-1 upregulation preceded the induction of inflammatory cytokines, a feature that deviated from model predictions. Model simulations suggested that direct induction of TfR1 after fungal recognition, independent of the Iron Regulatory Protein - Labile Iron Pool system, explains this finding. We anticipate that this model will contribute to a quantitative understanding of iron regulation as a fundamental host defense mechanism during aspergillosis.

**Importance:** Invasive pulmonary aspergillosis is a major cause of death among immunosuppressed individuals despite the best available therapy. Depriving the pathogen of iron is an essential component of host defense in this infection, but the mechanisms by which the host achieves this are complex. To understand how recruited macrophages mediate iron deprivation during the infection, we developed and validated a mechanistic computational model that integrates the available information in the field. The insights provided by this approach can help in designing iron modulation therapies as anti-fungal treatments.

## Introduction

The incidence of invasive aspergillosis continues to grow in tandem with the increasing use of immunosuppressive therapies (1, 2). Despite advances in diagnosis and therapy, mortality of invasive aspergillosis remains 30-60%, with most deaths occurring in patients on the best available therapy (3–5). The increasing prevalence of triazole resistance in this infection (6, 7) has raised the specter of a “perfect storm” due to a growing population of susceptible individuals with a diminished repertoire of treatment options (8).

Nutritional immunity, broadly defined as the restriction of essential nutrients from invading pathogens (9, 10), is an important component of antimicrobial host defenses (11, 12). The battle over iron represents the best-defined example of nutritional immunity (13–17), and is highly relevant to aspergillosis: Iron overload is an independent risk factor for invasive aspergillosis (18), and iron acquisition is essential to virulence of *Aspergillus* species (19, 20). The host sequestration of iron during the infection is implemented via multiple inter-related dynamic mechanisms, including cellular uptake of iron and heme, intracellular iron storage, and systemic suppression of iron availability, but the interplay of these host mechanisms during the infection is highly complex and poorly defined.

Mathematical modeling is a powerful tool for a principled integration of biological data and mechanisms, and the generation of novel hypotheses. While the response of activated macrophages to other lung infections has been studied using mathematical models (21–25), the published models do not address iron-related nutritional immunity in the setting of infection. Similarly, mathematical models of systemic iron regulation, macrophage iron-handling, and iron metabolism during erythropoiesis (26–28) are not specific to infections. Our group previously built a computational model of the competition of host immune cells and *Aspergillus* for access to iron during invasive infection (29), but this model did not include the intracellular handling of iron in macrophages in response to the infection. The focus of the current study is therefore to construct such a model and use it as a tool to integrate macrophage iron homeostasis upon contact with the fungus.

Here, we report the development of a novel mechanistic model of macrophage iron-handling during aspergillosis that quantitatively captures the molecular events in macrophage iron regulatory pathways over time. We generated a longitudinal transcriptomics data set from human monocyte-derived macrophages infected with *Aspergillus fumigatus* conidia and used it, together with available information from the literature, to construct a mathematical model that integrates pathogen recognition, transcriptional and post-transcriptional regulation, and autocrine/paracrine feedback loops influencing macrophage iron import, export, and storage. We validated this model with independent experimental data. The model was found to reproduce the dynamic changes in macrophage iron handling that were observed experimentally, which work in concert to limit the extracellular iron pool.

## Results

Transcriptomic analysis of macrophage-Aspergillus interaction shows activation of major innate immune pathways, including iron regulation.

We began with a time series experiment to measure the transcriptional landscape of human monocyte-derived macrophages that were incubated alone or with *A. fumigatus* conidia over 8h. We observed a time-dependent increase in the number of differentially expressed genes between macrophages incubated alone as compared to those co-cultured with *A. fumigatus*, beginning 4h after incubation (Figure 1A-B), consistent with the timing of shedding of the conidial rodlet layer (30). Principal component analysis of normalized read counts of the top 500 differentially expressed genes showed separation of infected and uninfected samples at 6 and 8h time points (Figure 1C).

**Figure 1.**
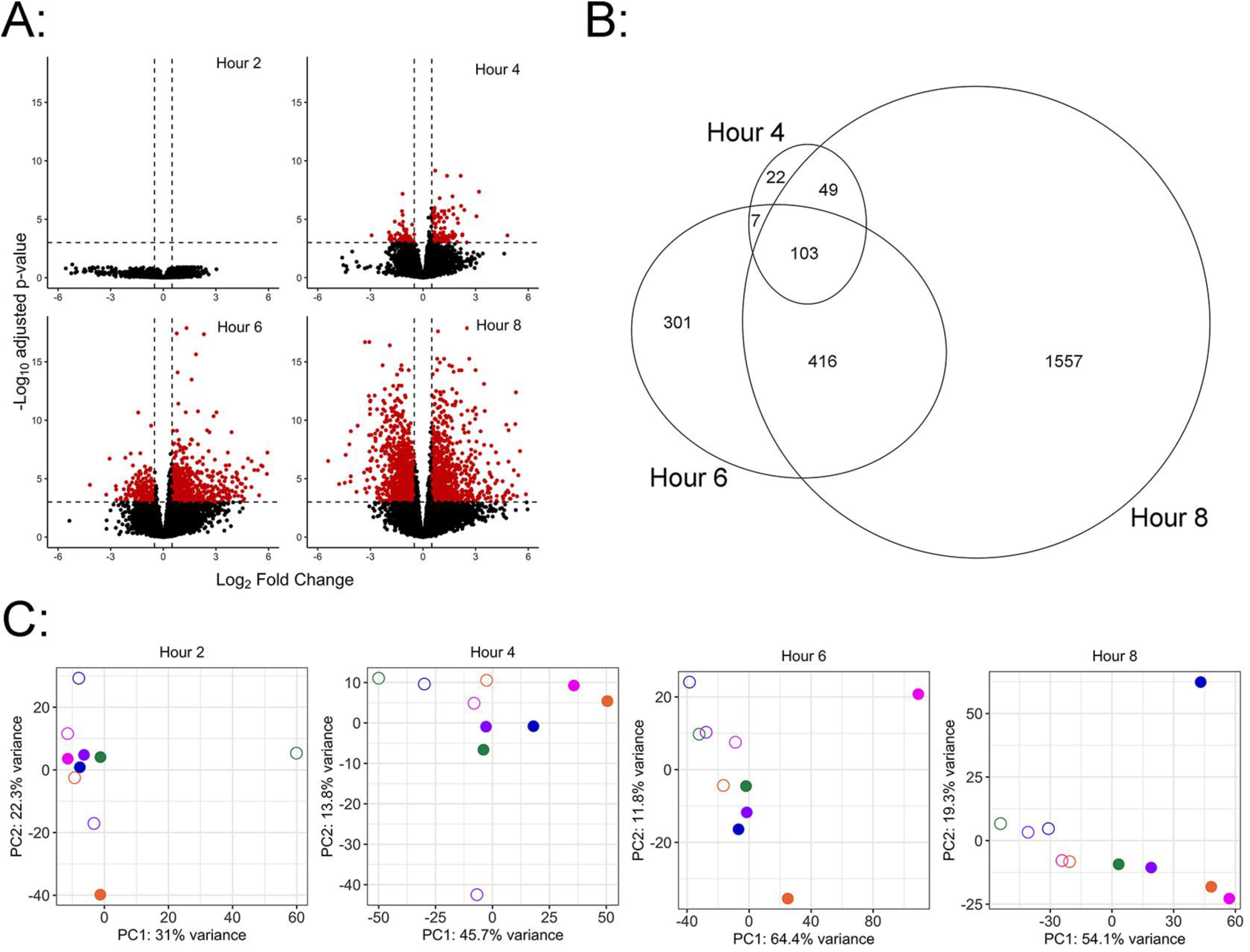
Differential expression analysis of macrophages infected with *Aspergillus*. A-B: Volcano plots and Euler diagram of genes with ≥2-fold differential expression in infected as compared to unifected macrophages with adjusted *p* value < 0.001. C: Principal component analysis plots of read counts of differentially expressed genes at each timepoint, after variance stabilizing transformation. Open and filled symbols indicate uninfected and infected cells, respectively, and the color of symbols denotes the donor.

We next performed enrichment analysis of differentially expressed genes at 4, 6, and 8h to obtain enriched Gene Ontology terms for biological processes, molecular functions and cellular components, and to identify related Reactome pathways. At 4h, we observed gene sets enriched for biological processes relating to phagocytosis (such as “phagosome acidification,” “endocytosis”, “regulation of intracellular pH”) and, notably, “transferrin transport” (Figure 2A). Consistent with these observations, “transferrin endocytosis and recycling” and “ROS and RNS production in phagocytosis” pathways were enriched for Reactome analysis at 4h (Figure 2B). Differential genes at 6 and 8h were enriched for Gene Ontology terms relating to multiple immune processes (e.g., “cellular response to lipopolysaccharides”, “response to tumor necrosis factor”, “cellular response to interleukin-1”, “neutrophil chemotaxis”, and “cell-cell signaling”; Figure 2B-C) and iron transport processes (eg., “transferrin transport” and “iron transport”). Reactome pathways analysis further confirmed activation of immune pathways with the enrichment of “NF-kappaB signaling pathway”, “TNF signaling pathway”, “cytokine-cytokine receptor pathway”, and “chemokine signaling pathway” at 6 and 8h (Figure 2E-F). We also observed that the “Iron uptake and transport” pathway was enriched at 8h, with 16 differentially regulated genes present in the pathway (Figure 2F).

**Figure 2.**
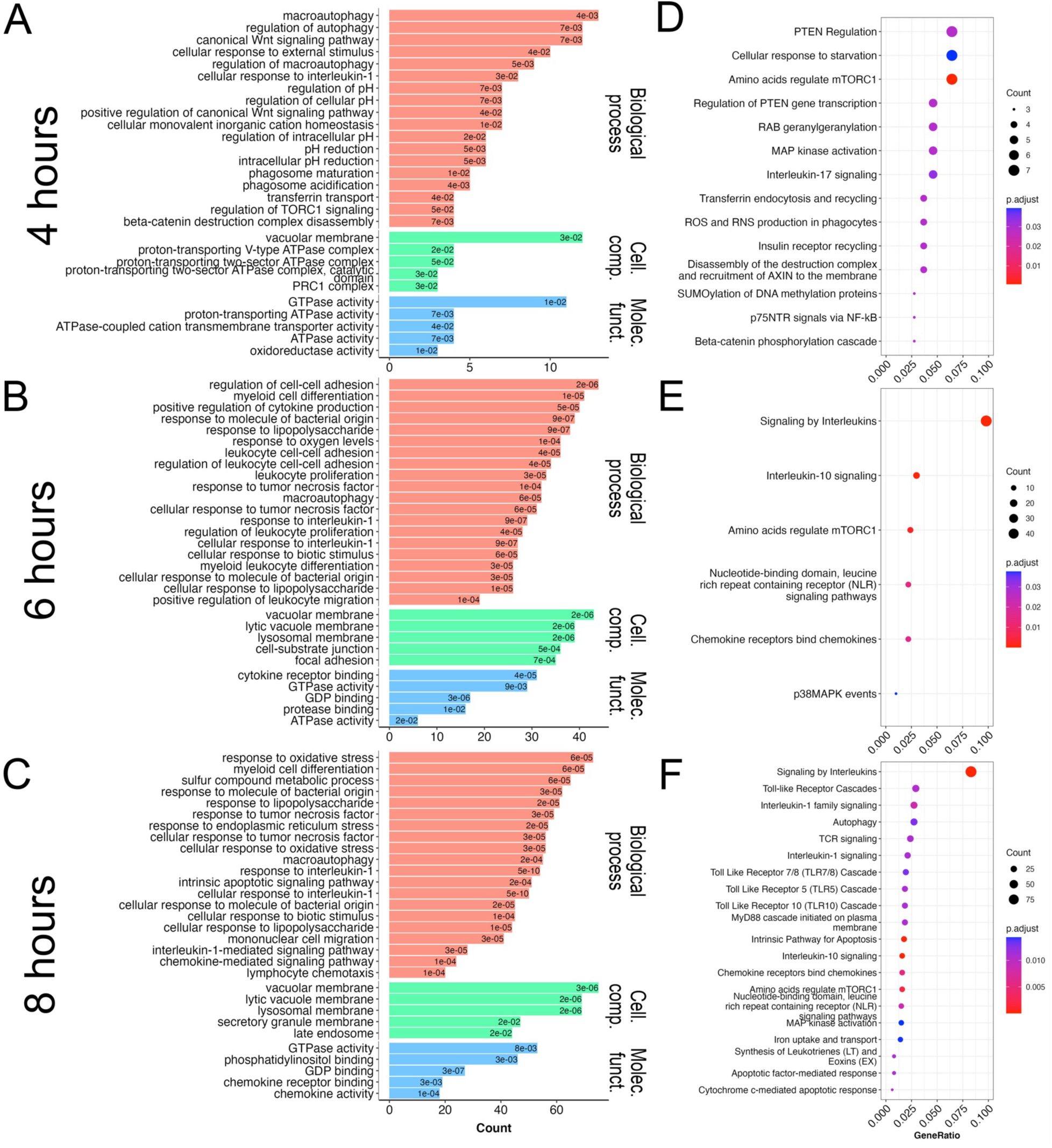
Enrichment analysis of differentially expressed genes in macrophages infected with *Aspergillus*. GO terms (A-C) and Reactome pathways (D-F) at 4, 6, and 8h after infection, respectively. Enrichment analysis was performed with differentially expressed genes (adjusted *p* value <0.001 and |Log_2_-fold change| ≥ 0.5). Enriched terms for Gene ontology (top 20 for biological processes, and top 5 for cellular components and molecular functions) and Reactome pathways (top 20) are reported. Cell. comp., cellular component; Molec. funct., molecular function.

We next focused on enrichment of iron-related genes. Using Gene Ontology and Reactome analyses, we observed pathways operational in iron regulation, including transferrin transport, iron transport, TNF- and IL-1-signaling. Ingenuity Pathway Analysis also indicated that, out of all “iron homeostasis” network molecules present in the IPA database, 26 genes involved in iron import/storage/export/transport pathways were differentially expressed. Unsupervised clustering of average expression data of iron-related genes obtained from AmiGO-2 revealed markedly different expression patterns for the 4, 6, and 8h infected culture groups, as compared to control and 2h infected samples (Figure 3). Overall, these data indicated early and robust activation of iron regulatory mechanisms in macrophages after fungal detection.

**Figure 3.**
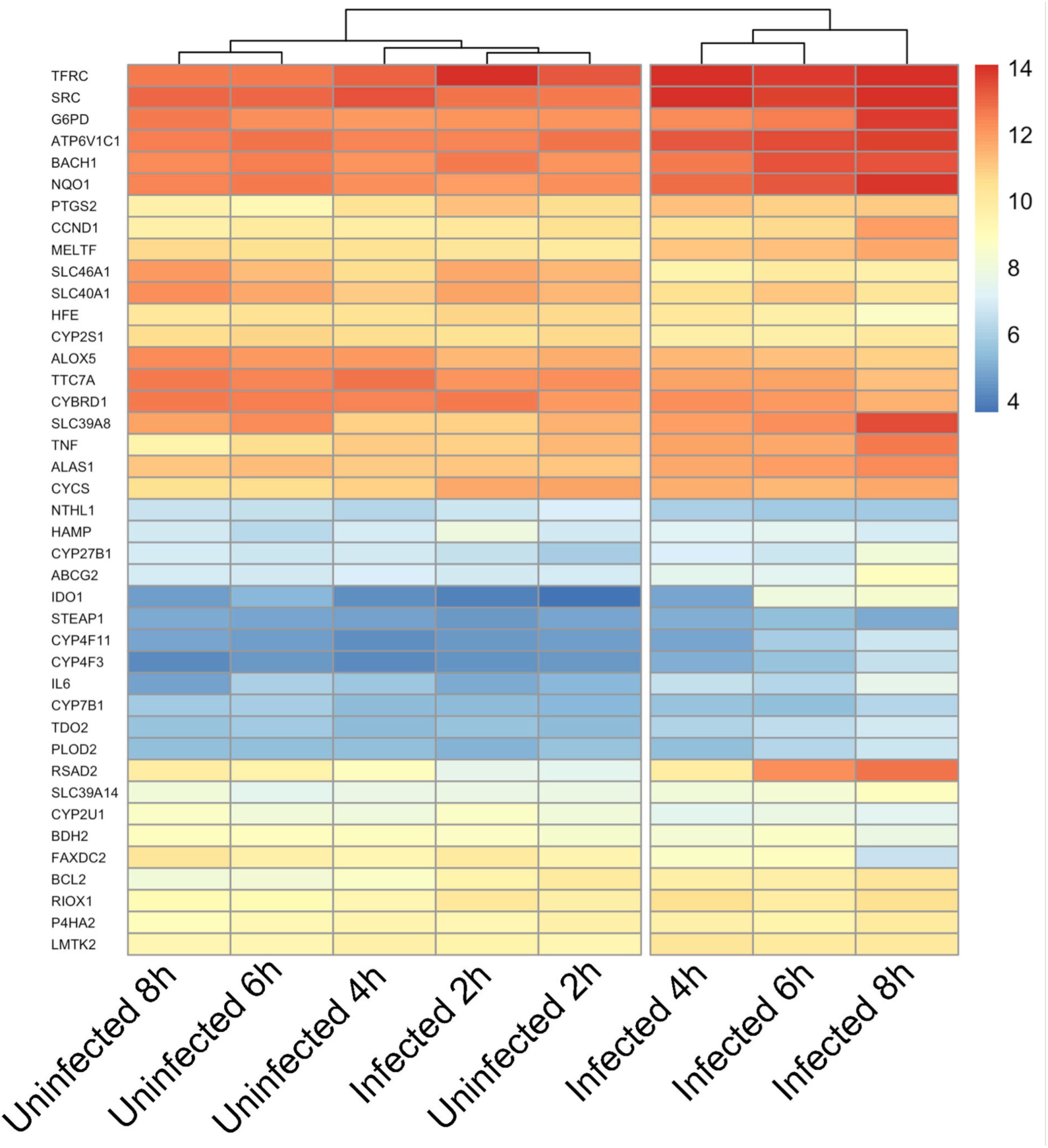
Heatmap of differentially regulated iron-associated genes after unsupervised clustering. Heatmap showing treatment groups on the x-axis and differentially regulated iron-associated genes with a |Log_2_-fold change| ≥ 1 on the y-axis. Each cell represents the median expression value of 5 biological replicates after variance stabilizing transformation on size factor normalized count data.

### Mathematical model of iron regulation in macrophages

We next integrated the RNA-seq results above with known biology to construct a mathematical model of iron regulation in monocyte-derived macrophages during an encounter with *Aspergillus* conidia. We reviewed the literature on each of the iron-related genes identified in our data and assessed their relevance to iron regulation and handling during fungal infections. We then established a set of molecules to include in the model (Table 1), based on our data and known literature (described below and depicted in Figure 4A). These molecules were incorporated into a static network (Figure 4B), which formed the basis for a discrete dynamical model, with each node in the model taking on three possible discrete states: 0, 1, 2. Hence, a model state is described as a vector of length 21 (the number of variables in the model), with entries 0, 1, or 2, representing different levels of each molecule. From a given initial state, the model evolves in discrete time steps by applying the regulatory rules in Table 2.

**Figure 4.**
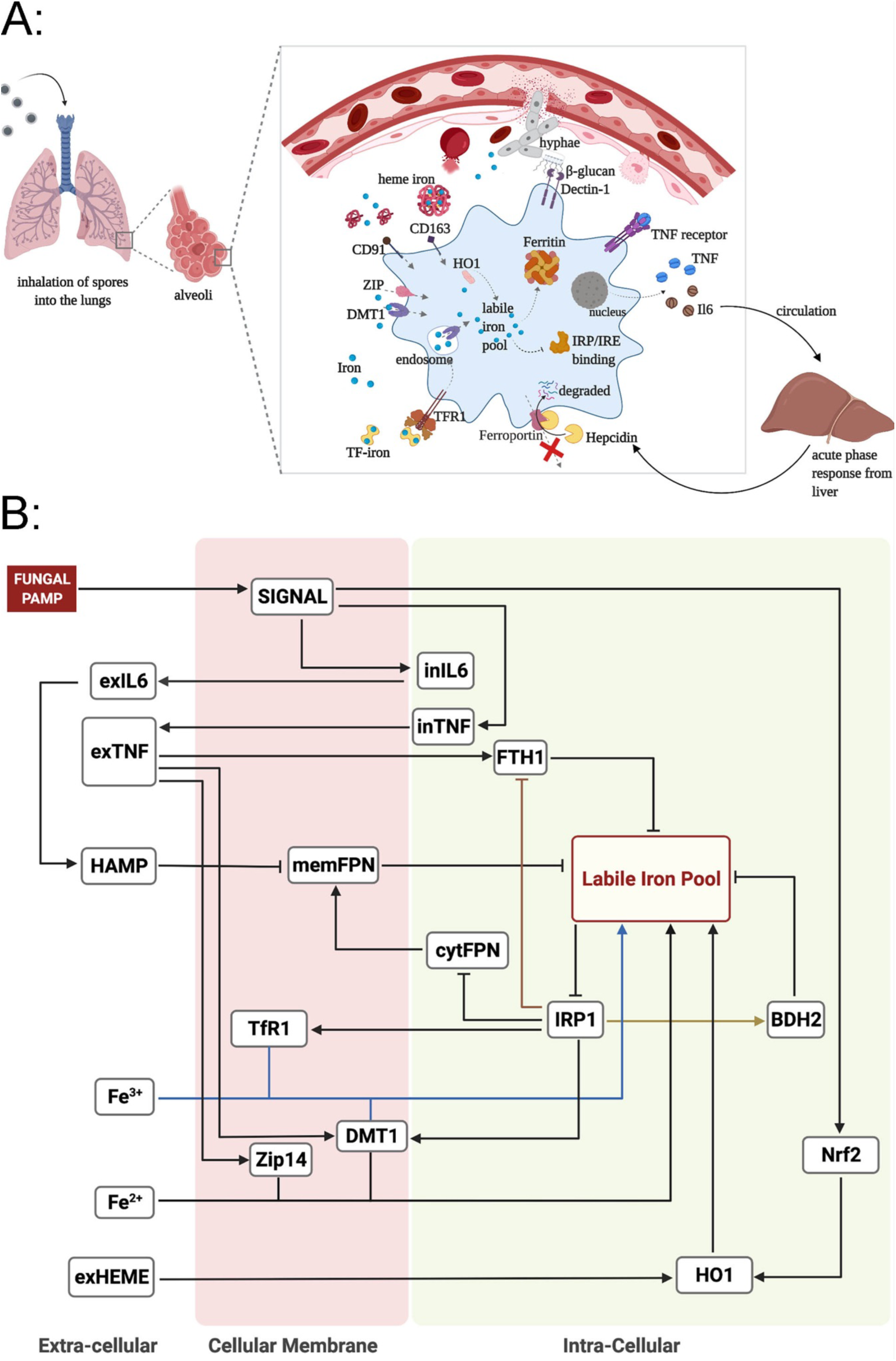
Computational model. A. Diagrammatic representation of key processes in iron regulation in macrophages during invasive pulmonary aspergillosis. B: Wiring diagram ofmacrophage iron regulation during invasive pulmonary aspergillosis. Pointed arrows represent activation and blunt arrows represent inhibition. Some arrows are colored for better visualization. Extracellular, membrane, cytoplasm and intracellular molecules are indicated by ex-, mem-, cyt- and in-prefixes. BDH2, 3-hydroxybutyrate dehydrogenase-2; DMT1, divalent metal transporter-1; Fe^2+^, ferrous iron forms; Fe^3+^, ferric iron forms; FPN, ferroportin; FTH1, ferritin heavy-chain-1; HAMP, hepcidin; HO1, heme oxygenase-1; IL6, interleukin-6; LIP, labile iron pool; IRP1, iron-regulatory protein-1; PAMP, pathogen-associated molecular pattern; TfR1, transferrin receptor-1; TNF, tumor necrosis factor; Zip14, zinc transporter-14. A-B, Created with BioRender.com.

**Table 1:** Biological description of variables and their possible states in the computational model. Extracellular, membrane, cytoplasm and intracellular molecules are indicated by ex-, mem-, cyt- and in-prefixes, respectively.

| Node | Name | Type | Location | Model States |  |  |
| --- | --- | --- | --- | --- | --- | --- |
|  |  |  |  | 0 | 1 | 2 |
| BDH2 | 3-hydroxybutyrate dehydrogenase-2 | Protein | Intracellular | Low expression | Normal | High expression |
| cytFPN | Cytoplasmic ferroportin | RNA | Intracellular | Low expression | Normal | High expression |
| DMT1 | Divalent metal transporter-1 | Protein, Importer | Membrane | Low activity | Normal | High activity |
| exIL6 | Interleukin-6 | Cytokine | Extracellular | Low expression | Normal | High expression |
| exHeme | Heme | Compound, Molecule | Extracellular | Low concentration | Normal | High concentration |
| exTNF | Tumor necrosis factor | Cytokine | Extracellular | Low expression | Normal | High expression |
| Fe<sup>2+</sup> | Labile ferrous iron ions | Ion | Extracellular | Low concentration | Normal | High concentration |
| Fe<sup>3+</sup> | Transferrin-bound ferric iron ions | Ion | Extracellular | Low concentration | Normal | High concentration |
| FTH1 | Ferritin heavy chain | Protein | Intracellular | Low expression | Normal | High expression |
| FUNGUS | *A. fumigatus* | Pathogen | Extracellular | Absent | Present | Present |
| HAMP | Hepcidin | Protein | Extracellular | Low concentration | Normal | High concentration |
| HO1 | Heme oxygenase-1 | Enzyme | Intracellular | Low expression | Normal | High expression |
| inIL6 | Interleukin-6 | Cytokine | Intracellular | Low expression | Normal | High expression |
| inTNF | Tumor necrosis factor | Cytokine | Intracellular | Low expression | Normal | High expression |
| IRP1 | Iron regulatory protein | Protein | Intracellular | Low activity | Normal | High activity |
| LIP | Labile iron pool | Molecules | Intracellular | Low concentration | Normal | High concentration |
| memFPN | Membrane-bound ferroportin | Protein, Exporter | Membrane | Low expression | Normal | High expression |
| Nrf2 | Nuclear factor erythroid factor 2-related factor 2 | Transcription factor | Intracellular | Low concentration | Normal | High concentration |
| TfR1 | Transferrin receptor-1 | Protein, Importer | Membrane | Low activity | Normal | High activity |
| SIGNAL | PAMP signaling after recognition of pathogen | Pathway | Intracellular | Low activity | High activity | High activity |
| Zip14 | Zrt- and Irt-like protein-14 | Protein, Importer | Membrane | Low activity | Normal | High activity |

**Table 2:** Update rules of model species and supporting literature citations. Continuity function accounts for the previous state of the target molecule when changing the state of the target molecule from a high to low level. Extracellular, membrane, cytoplasm and intracellular molecules are indicated by ex-, mem-, cyt- and in-prefixes, respectively. min, minimum; max, maximum; cont, continuity function.

| Target | Update Rules | Description |
| --- | --- | --- |
| BDH2 | IRP | BDH2 has an IRE motif on its 3' end. This interaction can lead to stabilization and increase in BDH2 (68). |
| cytFPN | cont(not(IRP1)) | IRP1 can bind to the IRE present on the 5' of ferroportin RNA inhibiting the translation of FPN (112). |
| DMT1 | max(cont(exTNF),<br>cont(IRP1)) | TNF induces the expression of DMT1 during infection, and DMT1 has an IRE element on 3'end of its mRNA (37, 113). |
| exIL6 | cont(inIL6) | IL6 is secreted into the extracellular environment (114). |
| exHeme | External Parameter |  |
| exTNF | cont(inTNF) | TNF is secreted into the extracellular environment (114). |
| Fe<sup>2+</sup> | External Parameter |  |
| Fe<sup>3+</sup> | External Parameter |  |
| FTH1 | max(cont(exTNF),<br>cont(not(IRP))) | TNF induces the expression of FTH1, and FTH1 has an IRE element on the 5' end of its mRNA for IRP regulation (37, 112, 115). |
| FUNGUS | Source Node |  |
| HAMP | cont(exIL6) | Hepcidin is produced by the |
|  |  | liver in response to the IL-6 (36, 87). |
| HO1 | min(exHEME, cont(Nrf2)) | Heme and NRF2 can activate expression of HO1 (116, 117). |
| inIL6 | SIGNAL | IL6 is produced in response to fungus (31–33, 118, 119). |
| inTNF | SIGNAL | TNF is produced in response to fungus (32, 33, 35, 118, 119). |
| IRP1 | cont(not(LIP)) | IRE-binding activity of IRP1 is high in iron-deplete conditions (120). |
| LIP | cont(min(max(min($\text{Fe}^{3+}$, TfR1), min($\text{Fe}^{2+}$, DMT1, Zip14), HO1), min(not(memFPN), not(BDH2), not(FTH1)))) | Import of transferrin-bound iron, free iron, and heme-iron increases intracellular iron. Storage of iron in ferritin, export of iron through ferroportin, and sequestration of iron by BDH2 decrease the labile iron pool in the cytosol (44). |
| memFPN | min(cont(cytFPN), not(HAMP)) | Translated ferroportin locates to cell membranes. Membrane ferroportin can be targeted by hepcidin for degradation (43). |
| Nrf2 | SIGNAL | NRF2 is produced in response to fungal beta-glucan (116). |
| TfR1 | max(cont(IRP1), SIGNAL) | IRP1 stabilizes TfR1 mRNA by binding to the IRE element on the 3' end of its mRNA thereby increasing total TfR1 (71, 121, 122). |
| SIGNAL | SIGNAL = low<br>if FUNGUS =0,<br>else SIGNAL = high | SIGNAL represents the activation of macrophages by the fungus (31). |
| Zip14 | cont(exTNF) | TNF induces expression of Zip14 (37). |

Macrophages recognize fungal pathogen-associated molecular patterns via surface pathogen-recognition receptors (31–34), leading to the production and secretion of TNF and IL-6 (33, 35). We represent this recognition by the presence of FUNGUS and activation of SIGNAL (Figure 4B). IL-6 induces the synthesis of hepcidin in the liver, a key iron regulatory hormone that is highly sensitive to systemic iron levels and, independently, inflammation (15, 36). TNF induces transcription of ferritin heavy chain-1 (FTH1), DMT1, and Zip14 via an autocrine/paracrine loop (37–41). Iron export occurs via the membrane protein ferroportin (42). Extracellular hepcidin binds to membrane ferroportin, mediating its internalization and subsequent proteolysis in endosomes, thereby lowering the efflux of iron to the extracellular environment (43, 44). We did not implement hepcidin production by macrophages in the model, because the effect has been reported to be comparatively negligible (45).

We incorporated three forms of extracellular iron import: Transferrin-bound iron (Fe^3+^), free labile iron (Fe^2+^), and heme-associated iron. The extracellular labile iron concentration will be exceedingly low (46), and is therefore not included. The concentrations of Fe^3+^, Fe^2+^, and heme-associated iron serve as external inputs, i.e., they are not regulated by other components of the model. Transferrin, the principal extracellular iron transport protein, binds ferric iron ions with high affinity and is internalized by receptor-mediated endocytosis via the transferrin receptor (47–50). Iron molecules then dissociate from transferrin and are shuttled into the cytosolic labile iron pool by endosomal membrane DMT1 (51). Similarly, labile iron can be taken up by the membrane proteins DMT1 or Zip14 from the extracellular environment and imported into the cytoplasm (40, 41, 52–55). With catabolism of hemoglobin (cell-free hemoglobin, resulting from infection-induced hemorrhage), there will be an increase in free iron and heme-bound iron (55–57). Free heme is complexed to hemopexin and taken up via CD91 (58, 59). We have represented this source of heme-associated iron as exHEME. Once imported, heme iron is converted to free iron by heme oxygenase-1 (HO1) and added to the labile iron pool, a redox-active form of iron present in the cytosolic environment (60, 61).

Excess iron in the cytosol is stored in ferritin, a 24-mer protein composed of light- and heavy-chain subunits (L- and H-ferritin, respectively) (62, 63). H-ferritin is also a ferroxidase enzyme, mediating the oxidation of ferrous to ferric iron for storage, and L-ferritin is important in the nucleation of ferric iron (64–66). We modeled FTH1 because it is transcriptionally regulated by inflammatory signals (39, 67). Cytosolic iron can also be bound by 2,5-dihydroxybenzoic acid (DHBA) molecules, also known as the mammalian siderophore (68, 69), represented in the model by BDH2, the enzyme that catalyzes the formation of 2,5-DHBA (69). The iron regulatory protein-1 regulates the intracellular labile iron pool by binding to iron responsive elements (IRE) of the untranslated 3’ and 5’ regions of mRNA of TfR1, DMT1, FPN, FTH1, and BDH2, thereby modulating the translation of iron storage, importer, and exporter proteins (68, 70–73): under low intracellular iron conditions, IRP1-IRE binding activity is high, inhibiting the translation of ferritins and FPN, and promoting the translation of TfR1, DMT1, and BDH2, with the opposite effect under iron-replete conditions (68, 71, 74).

### Computational model captures macrophage behavior during infection

We next used the computational model to simulate macrophage behavior under uninfected conditions. The external input parameters of the model were fixed to normal values of 1, and model dynamics were explored through complete enumeration of all state transitions. The first row of Figure 5 shows that the model reached a steady state, corresponding to a physiologically normal state. To test the model behavior under infection, we next simulated the presence of *A. fumigatus* (FUNGUS=Present). This resulted in macrophage-activated intracellular signaling pathways and the production of IL6, TNF, hepcidin, FTH1, Zip14, and DMT1. During infection, the model predicted low FPN transcription, membrane FPN, and the intracellular labile iron pool (LIP) in the steady state. Model simulation of an infected macrophage, in the presence of high iron, showed activation of HO1, in addition to the other iron regulatory molecules (Figure 5). HO1 catalyzes heme to ferrous iron, which adds to the intracellular labile iron pool (75–78), and is subsequently stored with ferritin.

**Figure 5.**
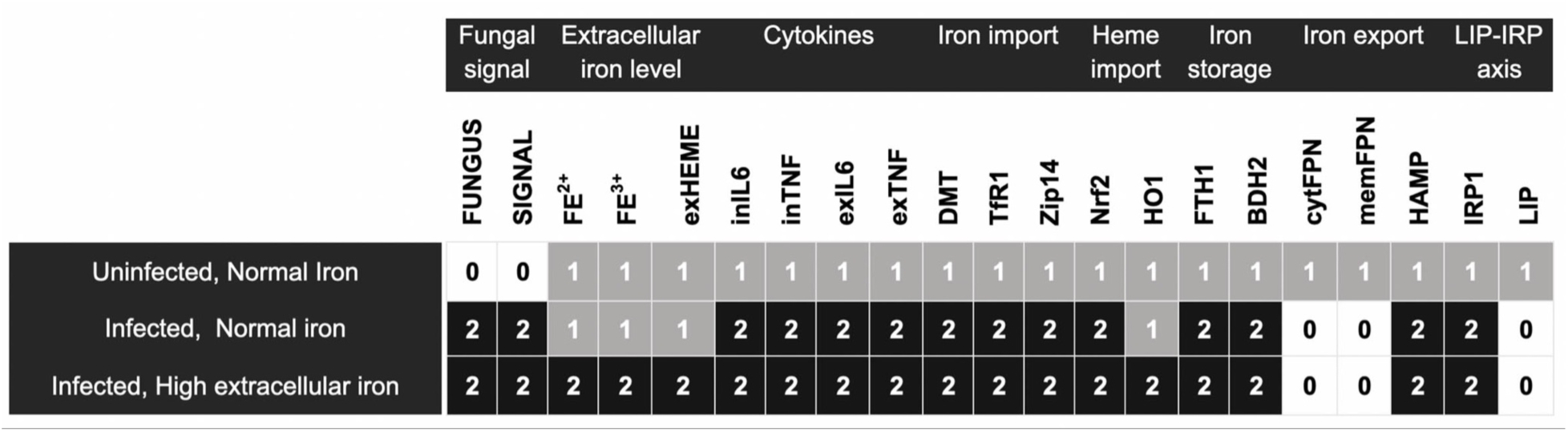
Different states of the computation model. A: Steady state simulations for the model under the conditions defined in Table 1 – uninfected macrophages in normal extracellular iron condition, infected macrophages in normal extracellular iron condition, and infected macrophages in high extracellular iron condition. 0, low; 1, medium/normal; 2, high.

### Validation of the computational model

Model simulation showed that, for each of the three conditions we consider, uninfected, infected, and infected with high iron levels, the model reaches a distinct steady state. We experimentally validated the model and its dynamic behavior in two ways. First, we used the temporal evolution as exhibited by the RNA-seq data set described above, which was not used for model construction. There, we had only used a differential expression analysis over the entire time course. We compared the steady states obtained from model simulation with the experimental data at 8h (Figure 6A), and second, we compared the model trajectory to the temporal dynamics of our longitudinal transcriptomic datasets (Figure 6B-D).

**Figure 6.**
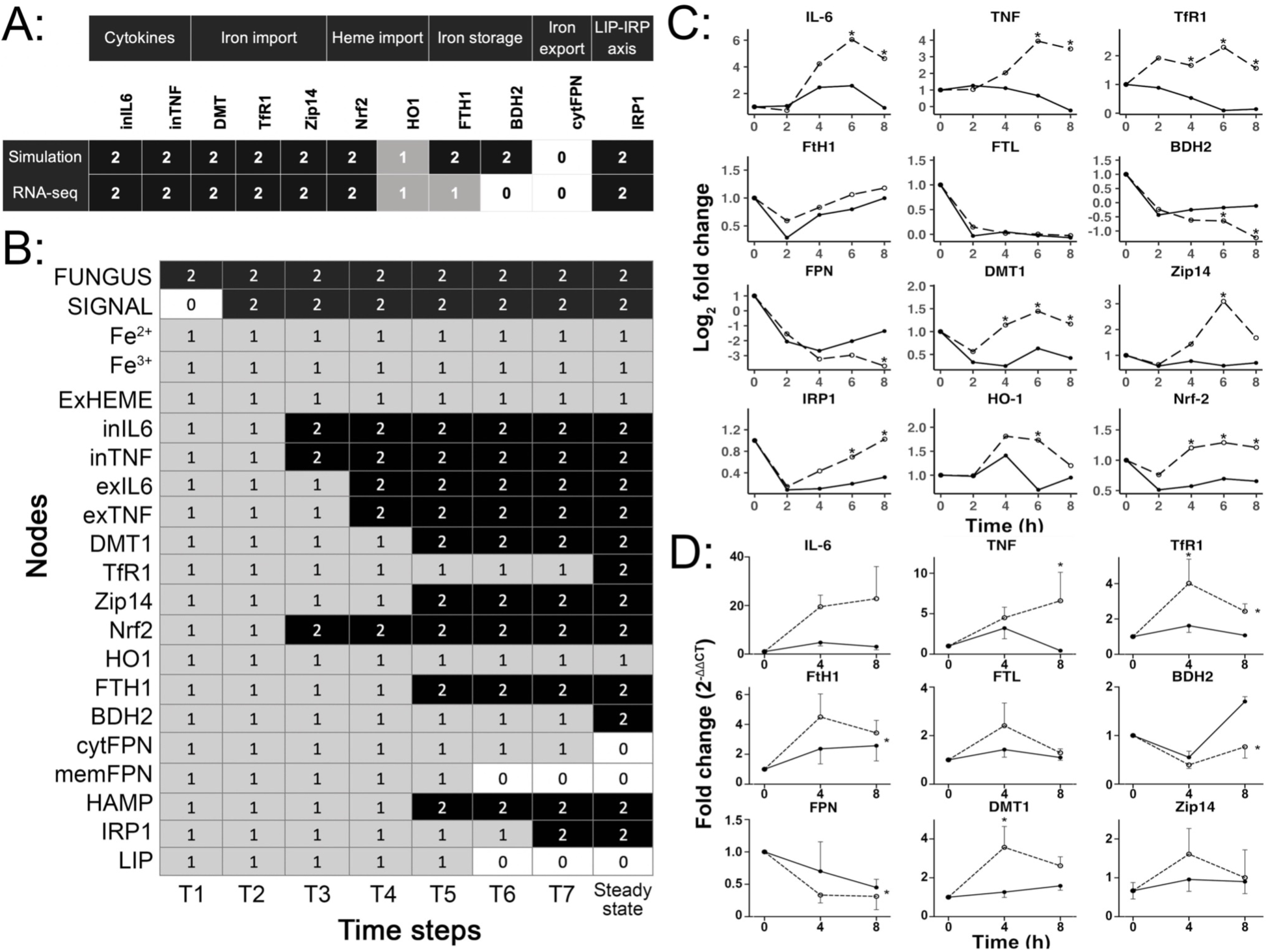
Validation of the computational model. A. Simulated steady states for infected macrophages under normal extracellular iron level and the RNA-seq data at 8h. Top row shows model output under conditions of exposure to the fungus and normal extracellular iron. Bottom row shows RNA-seq data discretized based on differential expression. B. Simulated time-series of the model output under conditions of exposure to the fungus and absent extracellular iron. C: RNA-seq experimental data was obtained from macrophage-*Aspergillus* co-cultures without external iron source. Read-counts were normalized by the library size and a value of 0.5 was added to the normalized counts to generate pseudo counts, which were then transformed with a Log_2_ scale. Log-scaled reads are plotted against time and actual raw read counts, and the line was fitted with loess regression. Counts were plotted using DESeq2 function plotCounts() method. *, *p* value < 0.05, and the line was fitted to the data with loess regression. D. Mean and SEM of qRT-PCR measurements from macrophages infected with *Aspergillus*. 0, down-regulated; 1, no change; 2, up-regulated. *, *p* value < 0.05; dotted lines, macrophage:fungus co-culture group; solid lines, control macrophage culture group.

The model includes 11 intracellular nodes that are measurable at the transcription level, namely TNF, IL6, Zip14, TfR1, DMT1, FTH1, BDH2, IRP1, HO1, NRF2, and FPN. We found that the model steady state matched the experimental data for all nodes, with the exception of BDH2 and FTH1 (Figure 6A). BDH2 simulation shows increased expression whereas the experimental data shows decreased expression, as previously reported in (79). This suggests that the model does not accurately capture BDH2 regulation, and that BDH2 is likely regulated by mechanisms independent of the known IRP regulation.

The model trajectory (Figure 6B) matched the experimental data from the RNA-seq dataset (Figure 6C) for 10 of 12 nodes: The induction of IL6 and NRF2 were observed in the simulation in earlier time steps and remained elevated until the steady state was reached, and both were differentially expressed from 4-8h in the experiment. Similarly, the iron importers DMT1 and Zip14 were activated at early stages in model simulations, and differentially upregulated from 4-8h in our experiment, suggesting induction of the iron import pathway post fungal recognition. Iron export, on the other hand, was inhibited upon the onset of infection in both simulation and in experimental data. The model indicated that membrane FPN is degraded at earlier time points whereas cytoplasmic FPN downregulation starts at later time steps in the simulation, suggesting that hepcidin regulation of FPN occurs prior to transcriptional regulation of FPN (Figure 6B). Concurring with the simulation data, measured FPN transcripts (cytFPN) also showed downregulation only at 8h in the experiment (Figure 6C). We also simulated molecules relevant to iron storage and chelation, ferritin and BDH2, respectively (Figure 6B). We observed a slight change in FTH1 expression with no statistical significance and no change in FTL expression (Figure 6C).

As a second step, experimental validation for these model predictions was performed on *Aspergillus*-infected macrophages using qRT-PCR (Figure 6D). These measurements matched the model trajectory (Figure 6B) for 8 out of 10 measured nodes and matched longitudinal transcriptome data (Figure 6C) for all nodes.

### Mathematical modeling suggests macrophage iron regulatory responses precede, and are independent of, generation of inflammatory cytokines

Mechanistic modeling can be used to assess whether known biology adequately accounts for observed behaviors. In our experimental data, we noted that the model accurately predicted the steady-state and temporal trajectory of TNF production, but while the model correctly predicted the steady-state TfR1 expression, it did not predict its trajectory correctly: In both the RNA-seq and qRT-PCR datasets, the upregulation of TfR1 occurred at 2-4h, preceding that of TNF at 8h, whereas the model predicted activation of TfR1 after the activation of IRP, indicating that the mechanisms incorporated into the model are not complete.

To assess this discordance, we tested different modifications of the model that would capture the trajectory of TfR1 expression matching the experimental observations. During fungal infections, nitric oxide has been reported to regulate TfR1, but our experimental data did not show expression of the iNOS gene, which is required for NO production in macrophages (80). We found that the only modification in the model that resulted in a steady state that reflected expected macrophage behavior during infection while maintaining all other trajectories and steady states was an additional regulation of TfR1 directly by the fungal node SIGNAL. With this modification, the model captures the observed phenotypic behavior in the experimental data. While TNF is one of the main cytokines operational in iron regulation during infection, early activation of TfR1, independent of IRP, suggests a separate activation pathway for enhancing iron import. The phenomenological regulation from SIGNAL to TfR1 suggests that further study of TfR1 regulation is needed to understand the mechanism of TfR1 induction during infection.

## Discussion

Multicellular hosts have evolved an evolutionarily ancient system of iron regulation in order to deprive invading pathogens of this essential nutrient: in response to diverse inflammatory and infectious stimuli, this system mediates iron sequestration within macrophages, a precipitous fall in plasma iron, plasma transferrin and transferrin saturation, and increase in plasma ferritin (10). Monocyte-derived macrophages recruited to the site of infection are at the crux of this system, controlling both the intra- and extra-cellular iron availability via modulation of iron import, storage, and export mechanisms, in response to a combination of signals from iron availability, pathogen recognition, inflammatory cytokines, and systemic hormones. These processes are intricately interdependent and are thus difficult to study in isolation, but can be integrated and understood using mechanistic computational modeling. We integrated the existing literature on macrophage iron control during fungal infection and transcriptional data obtained from co-culturing human monocyte-derived macrophages with fungal cells to develop such a model as related to aspergillosis.

The differential gene expression analysis of macrophages co-cultured with *A. fumigatus* revealed a transcriptional response that began between 2 and 4h after exposure to the fungus, consistent with prior work (81–84). Activated genes included those related to iron transport, storage, binding, and reductases, indicating the activation of iron regulatory mechanisms. In our experimental system, *Aspergillus* had access only to the small concentration of iron contained in the culture media and intracellular iron of killed macrophages. Analogously, inhaled *Aspergillus* conidia only have access to the iron-poor alveolar fluid and iron released from necrotic cells (85). But as the infection progresses, *Aspergillus* gains access to iron from tissue hemorrhage and hemoglobin catabolism in the host – a circumstance not represented in the experimental co-culture system, which models only the first 8h of the infection. Our model simulations were validated with experimental data and reflect the expected biology of *Aspergillus*-infected macrophages under different extracellular iron levels.

Macrophages can sequester iron via three mechanisms: increased iron import, increased iron storage, and decreased iron export. Regulation of iron export from macrophages (as well as duodenal enterocytes), mediated by the hepcidin-ferroportin axis, has been extensively documented in the literature in the context of normal iron homeostasis and during infections (86, 87). Our model shows that upregulation of iron importers in response to fungal detection plays an essential role in macrophage iron sequestration during infection, independent of iron export. The induction of transferrin receptor-1 and DMT1 in the experimental data validated this finding. In this context, iron regulation in macrophages was previously studied with a co-culture experiment of a RAW264.7 immortalized murine macrophage cell line (88). These cells showed an induction of ferritin, and a reduction of ferroportin expression after 7h of co-culture, but no change in TfR1 expression. The discrepancy between our findings and these results is likely due to differences between the murine cell line and primary human monocyte-derived macrophages, particularly the fact that RAW264.7 cells, derived from Balb/c mice, carry a homozygous non-functional NRAMP1 mutation, blocking iron shuttling between the phagosome and cytosol and impairing their iron homeostasis (89, 90).

Infection has been shown to result in reduction of extracellular iron to extremely low concentrations and in sequestration of iron inside macrophages (91–93). Our model simulation suggests that during infection, regardless of extracellular iron levels, the intracellular iron is stored by ferritin (Figure 5 and supplemental Figure 3). Our experimental data (Figure 6C-D), however, showed a small upregulation of FTH1 and no change in FTL. Of note, the baseline expression of FTH1 in both the infected and uninfected conditions was high, and we speculate that a slight upregulation in expression is enough to store a large quantity of iron, since one ferritin molecule can store up to 4500 iron ions (94). The model also elucidates the interplay between immune activation and the IRP-LIP axis of iron regulation: In Figure 5, the model indicated that normal iron-homeostasis under physiologic conditions is perturbed during infection, with high expression of DMT1, Zip14, TfR1, and ferritin mediated by inflammatory signals – IL6 and TNF – and independently of the IRP-LIP axis, resulting in augmented uptake and storage of iron in macrophages during infection. Consistent with this model prediction, the overriding influence of pro-inflammatory cytokines on iron regulation has been reported in other disease models. During inflammation in neuronal cells, the IL-6/JAK2/STAT3 pathway overrides iron homeostasis by dysregulation of hepcidin expression (95). In a study of the human monocytic cell line U937, TNF, IFN-ɑ, and IL-1β modulate iron metabolism by affecting macrophage iron uptake, TfR1 expression, intracellular iron handling, and ferritin mRNA levels (96).

Our computational model allowed us to capture the steady states of most model constituent molecules. Further analysis, however, indicated that the model did not capture some of the experimental observations. In particular, it did not capture the surprising biological observation of the activation of TfR1 at an earlier time point than TNF, both in the RNA-seq experiment and qRT-PCR. We modified the model to capture this feature through the inclusion of a hypothetical mechanism that activated TfR1 from a fungal signal. With this modification, the model agreed with the experimental data, suggesting a new hypothesis of TfR1 activation by an unknown molecule upstream of TNF. The transcriptional and post-transcriptional regulation of TfR1 by cellular iron deficiency and hypoxia, via the HIF-HRE and the IRP-IRE systems, is well-described (97), but its direct regulation by infectious stimuli has not, to our knowledge, been documented. Interestingly, TfR1 has been shown to localize to the early endosome of macrophages within minutes after phagocytosis of *Aspergillus* conidia (98), a timeline consistent with our experimental results and revised model predictions on the transcriptional activation of TfR1.

We recognize several limitations of our work: First, our computational model is based on data both from the literature and an *in vitro* co-culture system. It is likely that some aspects of macrophage behavior during the infection, such as extracellular signals relevant to iron regulation, are not captured by an *in vitro* experimental system. Second, the data we collected from the experimental co-culture system is limited to transcriptional changes, thus not capturing events such as protein phosphorylation or translocation. Both are partially addressed by supplementing the experimental data with extensive data from the literature. Third, the present work only pertains to the behavior of monocyte-derived macrophages recruited to the site of infection and does not capture the behavior of other cells (for example, alveolar macrophages and epithelial cells) relevant to iron handling during aspergillosis – and, as such, represents only part of the complex landscape of iron regulation during aspergillosis.

The current work has several implications for future studies, including suggesting several hypotheses for further exploration: First, as noted above, the mechanisms by which TfR1 is induced in macrophages independent of inflammatory cytokines, and the relevance of this induction to antifungal host defenses, are of interest. Second, the model simulation of BDH2 predicted an increased expression at the steady state due to its regulation by IRP, whereas experimental data showed BDH2 to be downregulated during infection (Figure 6C-D), suggesting that the known biology of regulation of BDH2 – and, by extrapolation, the role of the mammalian siderophore 2,5-DHBA – during fungal infection is incomplete. Third, the model that we have generated may be useful for predicting outcomes of iron-centered therapeutics, such as pharmacologic iron chelation (15, 99), by providing a better understanding of tissue macrophage iron handling during invasive aspergillosis. Finally, the current model can be incorporated into a multiscale computational model that incorporates the responses of other cell types with the fungus, synthesizing the available data in a systematic way and serving as an *in silico* laboratory.

## Materials and Methods

### Ethics statement

This study was conducted in accordance with the Declaration of Helsinki under a protocol approved by University of Florida Institutional Review Board.

### Fungal culture and harvest

*Aspergillus fumigatus* strain 13073 (American Type Culture Collection, Manassas, Virginia) was cultured on Sabouraud’s dextrose agar plates at 37°C for 14 days. Conidia were collected in PBS containing 0.1% Tween-80, filtered through sterile gauze, centrifuged at 700g, and resuspended in PBS, and concentration determined under a hemacytometer.

### Monocyte isolation, culture, and RNA extraction

Buffy coats from healthy volunteers (ages 21-78, 3 male and 3 female) were purchased (LifeSouth Community Blood Center, Gainesville, Florida). Mononuclear cells were isolated using Ficoll gradients and stored at -80°C degrees. CD14^+^ CD16^-^ monocytes were isolated using magnetic negative selection (EasySep Human Monocyte Isolation Kit, StemCell Technologies, Cambridge, Massachusetts) and differentiated into macrophages by culture in RPMI 1640 (Lonza, Morristown, NJ) supplemented with 2mM L-glutamine, 1mM Sodium Pyruvate Solution, 0.1mM nonessential amino-acids, 1% penicillin-streptomycin, 10% fetal bovine serum (Hyclone, Logan, UT), and 10ng/mL recombinant human macrophage colony-stimulating factor (Peprotech, East Windsor NJ) for 7 days. Flow cytometry was used to assess the purity of macrophages after 7 days of culture were assessed by flow cytometry (Supplemental Figure 1), according to a previously published protocol (100), using antibodies against CD68-BV711 (clone Y1/82A) and CD163-FITC (clone GHI/61), purchased from BD Biosciences (San Jose, California). Macrophages were co-cultured with *Aspergillus* conidia at a 1:1 ratio. At the beginning of the co-culture (time 0) and after 2, 4, 6, and 8h, cells were lysed (RLT buffer, Qiagen, Valencia, California) and RNA was extracted (RNeasy Plus mini kit, Qiagen) following the manufacturer’s instructions.

### RNA-seq library preparation, sequencing, and analysis

RNA-seq libraries were prepared and sequenced at the Jackson Laboratory for Genomic Medicine. Libraries were generated with KAPA-Stranded mRNA-seq kit (Roche Sequencing, Wilmington, Massachusetts) according to manufacturer’s instructions. Briefly, poly-A RNA was isolated from 300ng total RNA using oligo-dT magnetic beads. Purified RNA was then fragmented at 85°C for 6 mins, targeting fragments ranging 250-300bp. Fragmented RNA was reverse transcribed with an incubation of 25°C for 10 minutes, 42°C for 15 minutes and an inactivation step at 70°C for 15 minutes, followed by second strand synthesis at 16°C for 60mins. Double stranded cDNA fragments were purified using Ampure XP beads (Beckman Coulter Life Sciences, Indianapolis, Indiana). The dscDNA were then A-tailed, and ligated with Illumina unique adaptors (Illumina, San Diego, California). Adaptor-ligated DNA was purified using Ampure XP beads, 10 cycles of PCR amplification, and impurities were eliminated (Ampure XP beads, Beckman Coulter). RNA sequencing was performed on a HiSeq 4000 instrument (Illumina).

The sequenced raw RNA-seq reads were processed to generate read counts for alignment. In brief, the reads were checked for quality control using FASTQC v0.11.8 (www.bioinformatics.babraham.ac.uk/projects/fastqc/), and trimmed using Trimmomatic v0.39 using LEADING:3 TRAILING:3 SLIDINGWINDOW:4:15 MINLEN:36 and a predefined adapter list to be clipped from reads (101). FastQC and Trimmomatic were repeated until desired reads were obtained. MultiQC v1.7 was used to combine individual FastQC results for visualization (102). The trimmed reads were then aligned to the Ensembl GRCh38v96 reference genome using STAR v2.7.2b, and Qualimap v2.2.1 was used to check the quality of alignment (103, 104). Read-count matrix was created by using the column with reads for reverse strandedness (i.e. column 4) from readspergene.tab STAR output files. Samples from one donor were excluded from further analysis based on principal component analysis (PCA) performed on all samples, showing that this donor’s cells whether incubated alone or with conidia, formed a separate cluster from all other samples, with high biological variation (PC1 38%) relative to other donors and little response (PC2 20%) to the fungus (Supplemental Figure 2). PCA was performed on the top 500 variable genes in the datasets using the “prcomp()” function in R. Only genes expressed at 10 read counts or higher in at least 5 samples were processed further for differential expression analysis. DESeq 2 v1.24.0 was used to compute differentially expressed genes. The design matrix was created with 5 donors, 1 baseline, and 4 timepoints for control and co-culture groups each. Pairwise comparisons with Wald test (alpha=0.05) were performed for differential expression analysis. For computational validation of the model using RNA-seq data, RNA-seq read-counts were normalized by the library size and a value of 0.5 was added to the normalized counts to generate pseudo counts, which were then transformed with a Log_2_ scale. Log scaled reads are plotted against time and actual raw read counts, and the line was fitted with loess regression.

Gene Ontology analysis (105, 106) and Reactome (107) enrichment analysis was performed using the clusterProfiler (108, 109) package in R. These analyses were performed with top differentially expressed genes (adjusted *p* value <0.001 and |Log 2-Fold Change| >= 0.5) against *Homo sapiens* background separately for each time point. Gene Ontology terms and Reactome pathways at Benjamini-Hochberg-adjusted *p* values < 0.05 threshold were considered enriched. Ingenuity Pathway Analysis (www.qiagenbio-informatics.com/products/ingenuity-pathway-analysis) was performed to evaluate differentially expressed genes involved in major immune pathways in macrophages with the differentially expressed gene set (4h, 6h, 8h combined). To identify genes involved in iron regulation, we obtained “genes and gene predictions” from AmiGO2 (110) (amigo.geneontology.org/amigo), selecting “iron”.

### Quantitative reverse-transcription PCR

Macrophages were generated and co-incubated with *Aspergillus* as detailed above. After the incubation period, cells were suspended in RLT buffer (Qiagen, Valencia, CA), homogenized, passed through Qiashredder (Qiagen), and total RNA extracted using the RNAeasy mini kit (Qiagen) following the manufacturer’s instructions. Then, 1-0.2 μg of RNA was used to synthesize cDNA using the iScript cDNA synthesis kit (Bio-Rad, Hercules, CA). The cDNA template was mixed with iTAQ SYBR green universal super mix (Bio-Rad), and quantitative PCR was carried out on a CFX Connect system (Bio-Rad). Pre-designed human gene primers were purchased from Bio-Rad (supplemental Table). Human PPIA was amplified in parallel and used as the reference gene in quantification. Data are expressed as relative gene expression and were calculated using the 2^−ΔΔCT^ method.

### Mathematical model formulation and simulation

From the AmiGO2 database, known iron-related genes were extracted. Only differentially expressed iron genes from our transcriptional analysis from this list were taken into further consideration. A select few molecules that were not detected as differentially regulated in our transcriptional analysis but are reported as important in macrophage iron regulation in the literature were also considered. We reviewed literature on these molecules and built a static network depicting the relationship (regulatory edges of the network) between the molecules (nodes of the network) in Figure 4B. The static network is the basis for a time- and state-discrete dynamic model, with each node taking on three possible states: 0 (low), 1 (medium), 2 (high). We constructed transition functions encoding regulation of nodes and their evolution in discrete time steps (Table 2). Simulation code, in Python 3, is available at https://github.com/NutritionalLungImmunity/NLI_macrophage_iron_regulation. *<u>This model was our key discovery tool</u>*.

A possible model artifact is a variable change of more than one level per time step, e.g., from low to high without passing through medium. To avoid this, we applied a standard correction that forces this “continuity” property (111), which is known to not affect model features relevant to this study.

### Statistical analyses

Statistical analyses of RNA-seq data are described above. Other data were analyzed using the Prism software package (version 9.2.0, GraphPad Software, San Diego, California). The area-proportional Euler diagram was generated with EulerAPE (version 3.0.0; open source, http://www.eulerdiagrams.org/eulerAPE/). Comparisons of two groups over time or range of inocula was achieved using two-way ANOVA with Sidak multiple comparison test. A *p* value of <0.05 was considered statistically significant. In multiple comparison tests, multiplicity adjusted *p* values are reported.

---

**Supplemental Figure 1:**
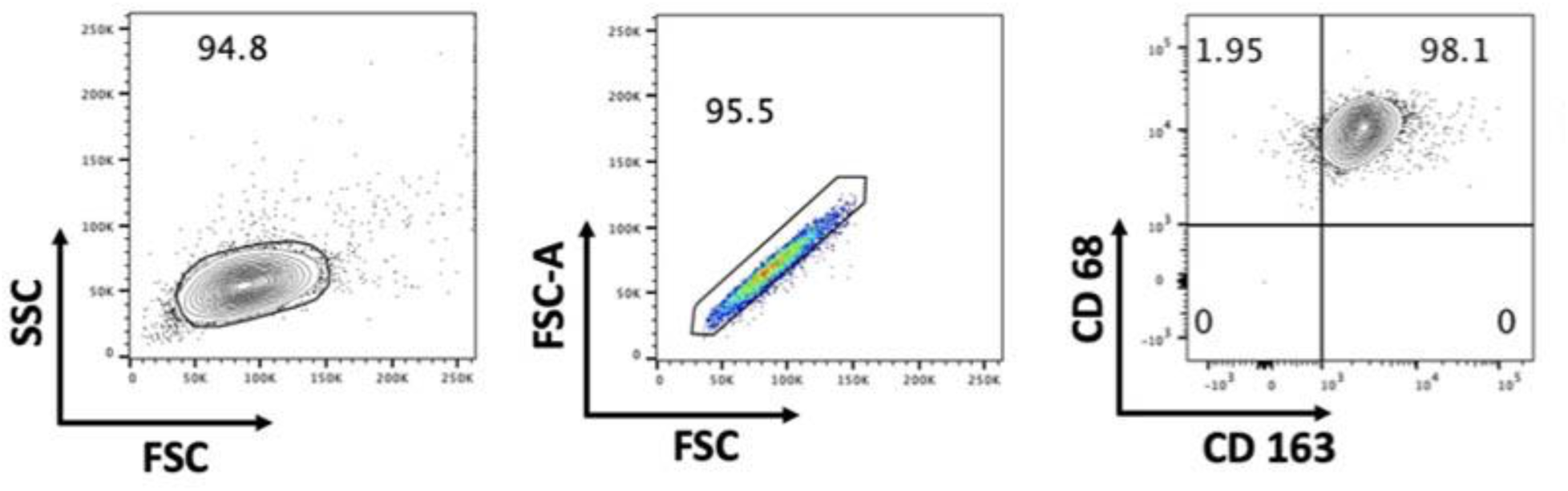
Representative flow cytometry plots showing purity of macrophages after culture for 7 days.

**Supplemental Figure 2:**
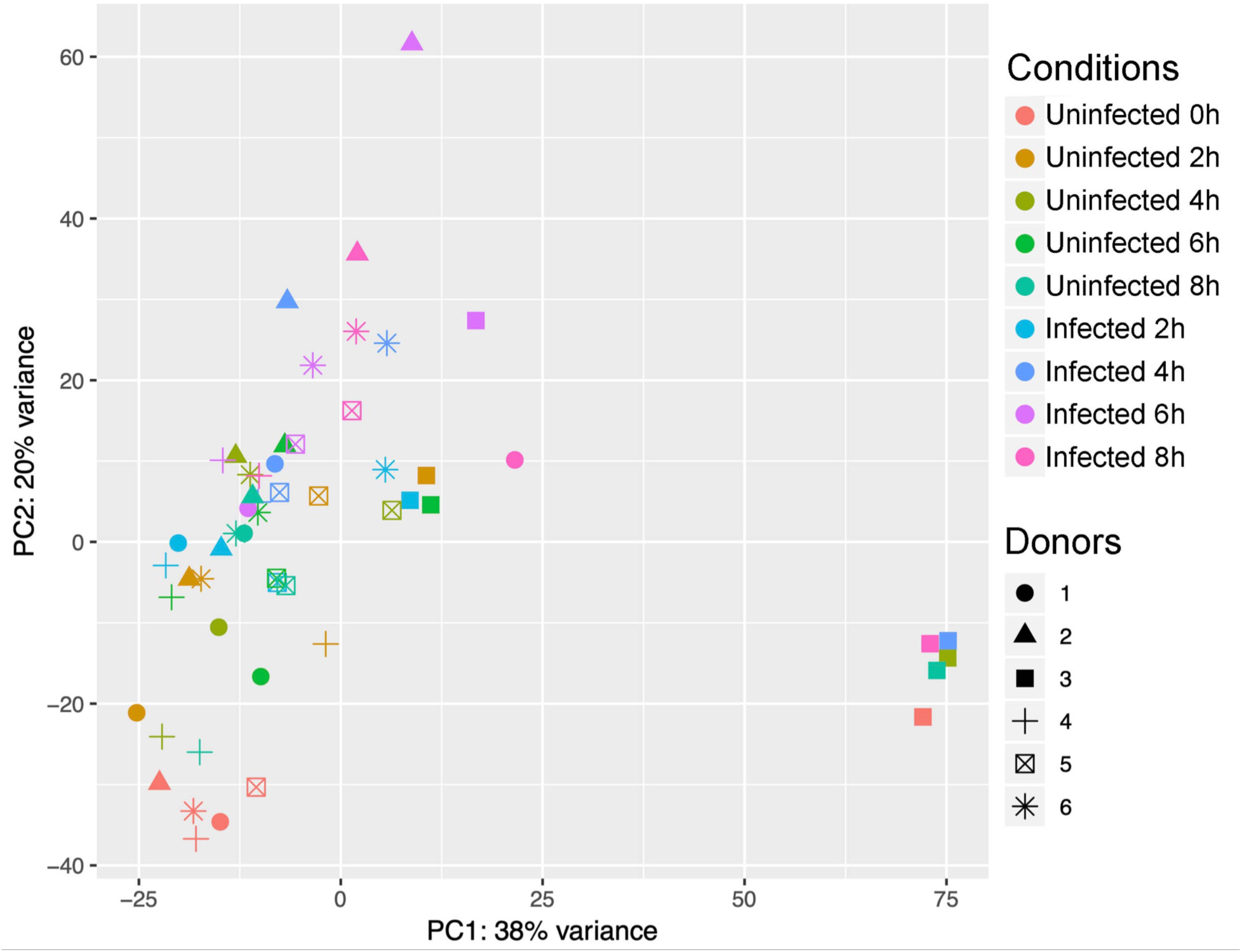
Principal component analysis plot of read counts after variance stabilizing transformation. Conditions are denoted by symbol color and donors by symbol type.

**Supplemental Figure 3:**
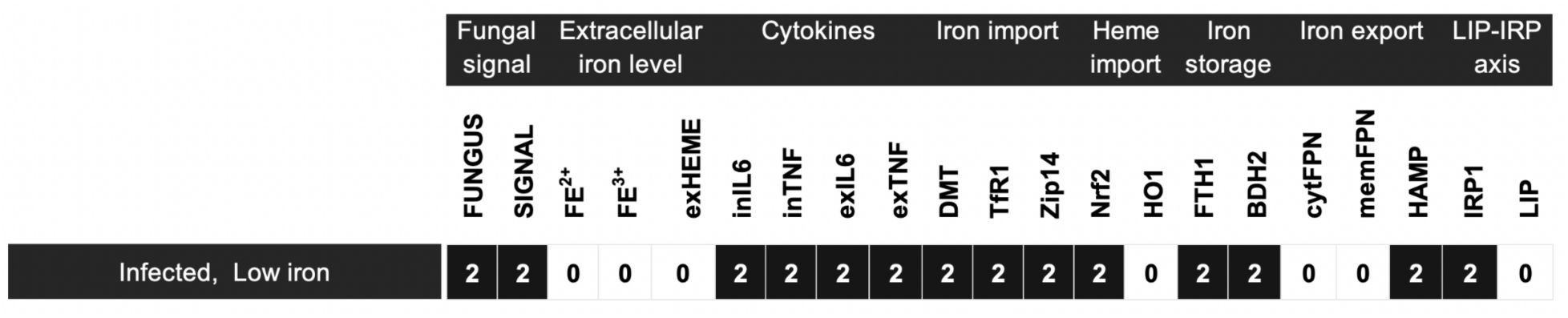
Simulated steady state of the model output under conditions of exposure to the fungus and absent extracellular iron.

**Supplemental Table:** PCR primer sequences used in the study.

| Gene | Unique Assay ID (BioRad Prime PCR) |
| --- | --- |
| BDH2 | qHSACED0046162 |
| FtH1 | qHsaCED0038139 |
| FTL | qHsaCED0057482 |
| IL-6 | qHsaCED0044677 |
| PPIA | qHsaCED0038620 |
| Slc11a2 | qHsaCID0012864 |
| Slc39a8 | qHsaCID0008870 |
| Slc39a14 | qHsaCED0002314 |
| Slc40a1 | qHsaCED0005662 |
| Tfr1 | qHsaCID0022106 |
| TNF | qHsaCED0037461 |

